# Modulation of negative affective biases in male rats may predict the rapid and sustained antidepressant effects of different NMDA antagonists

**DOI:** 10.1101/2025.02.26.640447

**Authors:** Justyna K Hinchcliffe, Katie Kamenish, Julia M. Bartlett, Roberto Arban, Bastian Hengerer, Emma S.J. Robinson

## Abstract

Affective biases influence cognitive and emotional behaviour and play a significant role in major depressive disorder. We have shown that the NMDA antagonist and rapid-acting antidepressant, ketamine, selectively modulates affective biases, a neuropsychological effect which may underlie its efficacy. Clinical studies with different NMDA antagonists have found mixed effects but the reasons for these differences in efficacy are not understood. This study used a rat model of negative affective biases to investigate if different NMDA antagonists also differentially modulated affective biases. Dose-response experiments using the NMDA antagonists lanicemine, memantine, CP101,606, phencyclidine (PCP) and ephenidine, and ketamine metabolite, (2R,6R)-hydroxynorketamine (HNK), were performed to test their acute and sustained (24hrs) effects, and specificity of affective bias modulation. Our results showed that HNK, PCP, lanicemine, and ephenidine acutely attenuate negative biases. HNK, CP-101,606, and ephenidine’s effects were sustained at 24hrs post-treatment with a positive bias observed for HNK and a tendency towards a positive affective bias for CP101,606 and ephenidine. Lanicemine’s effects were sustained at 24hrs but only at the highest dose and PCP and memantine had no effects. Considering these findings in the context of clinical observations, we suggest that the ability to induce a sustained modulation of affective biases corresponds with therapeutic effects. The differences in efficacy observed with NMDA antagonists may be related to their ion trapping properties and those with very high and low ion trapping properties are less effective than those with more moderate ion trapping effects such as ketamine or ephenidine or the subunit selective antagonist, CP101,606.

## Introduction

The term affective bias or cognitive affective bias is used to describe how affective states influence information processing and are thought to play an important role in major depressive disorder (MDD) (1). Different cognitive domains have been shown to be modulated by positive or negative affective states including learning and memory, decision-making and attention (2, 3). These affective biases can change perception, memory retrieval and decision-making, and in MDD amplifying negative emotions and thoughts, while diminishing positive ones (2, 3). Negative affective biases are hypothesised to play an important role in both the development and perpetuation of mood disorders (1, 2, 4) and modulation of negative affective biases has been shown with conventional delayed onset antidepressants in both humans (1) and rats (5-7), and rapid acting antidepressants (RAADs) in rats (7, 8). Exploring this neuropsychological hypothesis of antidepressant efficacy in healthy volunteers and patients, studies find a dissociation between the acute neuropsychological effects and the delayed subjective self-reported effects (1, 4). Recent findings in a rodent model of negative affective biases suggests that the NMDA antagonist, ketamine and other RAADs, such as the psychedelic, psilocybin, also interact with negative affective biases but through mechanism which are distinct from those seen with conventional antidepressants (8). The ability of these RAADs to modulate affective biases associated with past experiences has been hypothesised to explain their rapid and sustained antidepressant effects (8).

Since the first clinical study reporting a rapid and sustained antidepressant effect with ketamine (9), both clinical trials and studies in rodent models of depression have explored the potential efficacy of other NMDA antagonists. Ketamine’s clinical benefits in MDD are observed within hours of acute administration and last for up to 14 days (9, 10). However, studies with other NMDA antagonists, and even different doses of ketamine, have found mixed results (10). Different hypotheses have been proposed to explain this lack of efficacy including ketamine acting through non-NMDA mechanisms e.g. opioid receptors or via its active metabolite, hydroxynorketamine (11, 12). NMDA receptors are complex and these ligand-gated ion changes can be formed from different sub-unit (NR2A, 2B, 2C and 2D) compositions with different regional and cellular distributions which influences their functional effects (13). The pharmacological modulation of NMDA receptors includes targeting the ligand-binding domain and modulatory sites but most commonly, the channel pore. Ketamine is a non-competitive channel blocker with moderate to high ion trapping properties. Other non-competitive NMDA antagonists exist which have lower ion trapping properties e.g. lanicemine, both low affinity and low ion trapping properties e.g. memantine, or very high ion trapping properties e.g. PCP.

In this study we used a rodent model of affective biases to investigate the effects of different NMDA antagonists as well as the ketamine metabolite, HNK. We have previously shown that low but not high dose ketamine selectively modulates affective biases in the rat affective bias test (8). We also found ketamine and the NR2B selective antagonists CP101,606, modulate biases in decision-making (14). We localised these effects to the medial prefrontal cortex (8) and in the ABT, we show that the sustained effects of ketamine involve experience-dependent plasticity with negatively biased memories re-learnt post-treatment with a more positive affective valence. To further investigate the mechanisms underlying ketamine’s affective bias modulation and compare different NMDA antagonists we use our two ABT protocols. To investigate the acute versus sustained modulation of affective biases, we first induce a negatively biased memory then administer the treatment either 1hr or 24hrs before testing memory retrieval (15). We also tested any treatments which induced a significant acute modulation of affective biases using our control reward learning assay so we can determine if any effects observed were specific. We first tested the effects of the ketamine metabolite (2*R*,6*R*)-hydroxynorketamine which has previously been reported to induce antidepressant-like effects in preclinical models (12, 16). We next compared the effects of two NMDA antagonists which have failed in human clinical trials for MDD, lanicemine, low ion trapping, and memantine, low affinity and moderate ion trapping, and the high ion trapping NMDA antagonist PCP. We also tested ephenidine (two ringed *N*-ethyl-1,2-diphenylethylamine) which has similar pharmacological effects at the NMDA receptor as ketamine but lacks affinity for opioid receptors and does not produce the metabolite HNK. The final compound tested was CP101,606, the GluN2B subunit selective NMDAR antagonist that has demonstrated promising rapid acting antidepressant effects in human clinical trials (17) but failed to progress due to toxicity issues.

### Animals

Seven separate cohorts of male Lister Hooded rats (Envigo, UK) were used in these experiments (n=10-12 per group; Table S1). This study utilized only male rats; however, previous research indicates that similar affective biases are observed in both male and female rats (5). The sample size was based on our previous affective bias test studies and a meta-analysis which suggested a medium to large effect size for the drug-induced negative bias and reward-induced bias in Lister Hooded rats (5, 18). All animals were pair-housed in standard enriched laboratory cages (55×35x21cm) with woodchip, paper bedding, cotton rope, wood chew, cardboard tube and red Perspex house (30×17×10cm), under a 12:12h reverse light-dark cycle (lights off at 08:00h) and in temperature-controlled conditions (21±1°C). Rats were food restricted to approximately 90% of their free-feeding weights matched to the normal growth curve [∼18 g of food per rat/day laboratory chow (Purina, UK)] and were provided with water *ad libitum*. The behavioural procedures and testing were performed during the animals’ active phase between 09:00h and 17:00h. All experimental procedures were conducted under the UK Animals (Scientific Procedures) Act 1986 and were approved by the University of Bristol Animal Welfare and Ethical Review Body and UK Home Office (PPL 9516065).

### Affective Bias Test training and testing

The testing apparatus and pre-training handling and training protocol followed that of Hinchcliffe & Robinson, 2024 (15). Training involved animals learning to dig in ceramic bowls containing sawdust over 5 days with increasing levels of difficulty. The final session tested their ability to learn the task rule in a novel discrimination session where rats had to dig in the correct digging substrate to finding a food reward. Choice of the reward-paired substrate was marked as a ‘correct’ trial, digging in the unrewarded substrate was classified as an ‘incorrect’ trial and if an animal failed to approach and explore the bowls within 10 seconds, the trial was recorded to be an ‘omission’. Trials were continued until the rat achieved six consecutive correct choices for the reward-paired substrate. The discrimination session was used to confirm that the animals could achieve our learning criterion of six consecutive correct trials in less than 20 trials. Once animals successfully reached criteria in the discrimination session, they were considered trained. All animals then progressed to a reward learning assay protocol to confirm that they would exhibit a reward-induced bias and were therefore performing the task correctly and making their choice based on the memory associated with the digging substrate.

During testing, each week consisted of four consecutive pairing sessions to generate two independent cue-specific experiences. Using a within-subject design, each animal learnt a specific substrate-reward association under either a control or treatment conditions. A choice test on the fifth (acute modulation or reward learning assay) or sixth (sustained modulation) day of the same week assessed memory retrieval with or without drug pre-treatment. Details of all pairing sessions and choice test procedures are explained in the supplementary methods (Tables S2A and S2B). All drug treatments, pairing substrates and order of presentation were fully randomised in all studies. Affective biases generated using this protocol were quantified during the choice test when the two previously rewarded substrates (‘A’ and ‘B’) were presented at the same time over 30 randomly reinforced trials (for details, see (15)). The animals’ choices and latency to dig were recorded.

### Drugs

The drugs used to induce a negative affective bias in rats were corticosterone (10mg/kg, SC) and FG7142, benzodiazepine inverse agonist (3mg/kg, SC). The NMDA receptor antagonists tested were HNK (0.3, 1, 3mg/kg, IP), PCP (0.1, 0.3, 1mg/kg, IP), memantine (0.3, 1, 3mg/kg, IP), lanicemine (1, 3, 10mg/kg, IP), CP101,606 (1, 3mg/kg, IP) and ephenidine (1mg/kg, IP) (Table S1). All drugs were purchased from Merck (previously Sigma-Aldrich) or MedChemExpress. Ephenidine was kindly provided by Dr Jason Wallach. CP101,606 was dissolved in vehicle of 5% DMSO, 10% cremophor, 85% saline and all remaining NMDA antagonists were dissolved in saline vehicle in a dose volume 1ml/kg. For corticosterone or FG7142-induced negative affective biases, we selected a dose previously shown to induce a robust negative affective bias in the affective bias test (7, 8). PCP, memantine, lanicemine and CP101,606 and doses were based on previous studies using judgement bias task (14). HNK doses were based on previous publications (11, 16). Ephenidine dose was chosen based on its pharmacodynamic profile (19). Intraperitoneal injection procedures were done using a low-stress, non-restraint method developed in our research group (20, 21). All animals were habituated to the position required for dosing for five days prior to the experiments. All subcutaneous injections were performed with minimal animal restraint and injected on their left or right flank (changing daily). In all experiments, a within-subject design was used, with the experimenter blind to treatment and with a fully counterbalanced experimental design.

### Data analysis

Data were analysed and figures were created using GraphPad Prism 10.2.0 (GraphPad Software, USA). Choice bias score was calculated as the number of choices made for the drug-paired substrate (affective bias test) or two pellets-paired substrate (reward learning assay) divided by the total number of trials multiplied by 100 to give a percentage. A value of 50 was then subtracted to give a score where a choice bias towards the drug-paired substrate gave a positive value and a bias towards the control-paired substrate gave a negative value. For the memory retrieval studies involving a FG7142 or corticosterone-induced negative bias, animals that did not exhibit the expected negative bias under vehicle treatment were excluded. Values that were more than 2 standard deviations from the group mean were also excluded. Choice bias scores and response latency scores during the choice test were analysed using a repeated measures ANOVA with treatment as the within-subject factor, and as a post-hoc analysis pairwise comparisons were made using a two-tailed paired t-test or Dunnett’s test depending on the number of group comparisons. Individual positive or negative affective biases were also analysed using a one-sample t-test against a null hypothesis mean of 0% choice bias. For each animal, mean trials to criterion and latency to dig during the affective bias test pairing sessions and choice test were analysed using a repeated measures ANOVA with treatment as the factor or a two-tailed paired t-test, with post-hoc pairwise comparisons made using a two-tailed paired t-test comparison between control (vehicle/low reward:1 pellet) and treatment/manipulation (corticosterone/FG7142/high reward: 2 pellets) for each week (drug-induced negative bias retrieval studies and reward learning assay). Analysis of the choice latency and trials to criterion was made to determine the presence of any non-specific effects of treatment, such as sedation. A Shapiro-Wilk test was used to determine a normal distribution for the % choice bias, trials to criterion, and mean latency to dig during pairing sessions and choice test. Mauchly’s sphericity test was used to validate a repeated measures ANOVA.

## Results

To investigate the modulation of negatively biased memory, we initially induced a negative affective bias using either the FG7142 (3mg/kg) or corticosterone (10mg/kg). Following this, we administered the test compound either 1 hour or 24 hours before conducting the choice test. To determine if the effects of treatment were specific to an affective state-induced bias, we utilised a reward learning assay as a control for general memory deficits. In all studies, the control rats made fewer choices for the treatment-paired (either FG7142 or corticosterone-paired) digging substrate consistent with a negatively biased memory (one-sample t-test, p<0.0001, Fig. 1A-F).

**Figure 1:**
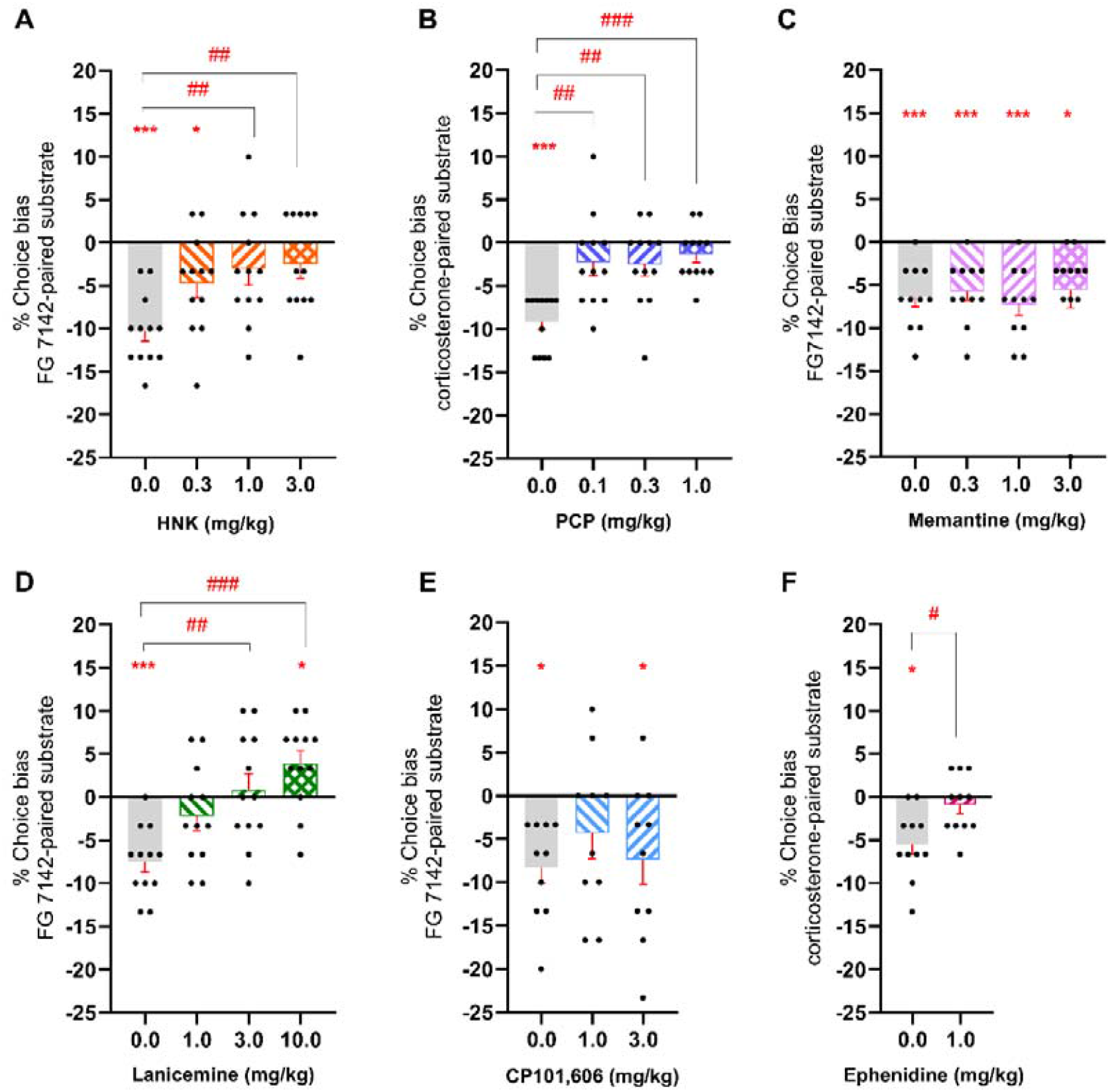
HNK, PCP, lanicemine and ephenidine attenuate negative affective bias in rats. Following induction of a negative affective bias with FG7142 or corticosterone, male rats were injected with HNK (0.3, 1.0, 3.0mg/kg; t=-20min; n=12; panel A), PCP (0.1, 0.3, 1.0mg/kg; t=-60min; n=12; panel B), memantine (0.3, 1.0, 3.0mg/kg; t=-60min; n=11; panel C), lanicemine (1.0, 3.0, 10.0mg/kg; t=-60min; n=12, panel D), CP101,606 (1.0, 3.0mg/kg; t=-60min; n=11, panel E) or ephenidine (1.0mg/kg; t=-60min; n=11, panel F) prior to choice test. Only two highest doses of HNK 1.0 and 3.0mg/kg, all doses of PCP, the middle dose of lanicemine 3.0mg/kg and one dose of ephenidine 1.0mg/kg significantly attenuated previously learnt negatively biased memories in rats. The highest dose of lanicemine not only attenuated the negative bias but rats also shown a positive bias at the group level (panel D). Data are shown as mean % choice bias ± SEM (bars) as well as individual data points (dots, n=11-12). Data were analyzed with one sample t-test against a null hypothesis mean of 0% choice bias (*p<0.05, ***p<0.001) and pairwise comparisons were done using Dunnett’s test following main effect of treatment in RM ANOVA (##p<0.01, ###p<0.001).

Acute HNK at 1.0 and 3.0mg/kg (repeated measures ANOVA, F3, 33=4.861, p=0.0066, n=12, with Dunnett’s test, p<0.001, Fig. 1A), and PCP at all doses (repeated measures ANOVA, F3, 33=7.332, p=0.0007, n=12, with Dunnett’s test, p<0.001, Fig. 1B), attenuated the negative bias when administered 1hr prior to the choice test,. The negative bias was also attenuated when animals received the two highest doses 3.0 and 10.0mg/kg of lanicemine (repeated measures ANOVA, F3,33=8.316, p=0.0003, n=12, with Dunnett’s test, p<0.001, Fig. 1D) and 1.0mg/kg ephenidine (two-tailed paired t-test: t10=3.012, p=0.0131 vs vehicle, n=11, Fig. 1F). The highest dose of lanicemine not only attenuated the negative bias but rats also showed a positive bias at the group level (one-sample t-test, p=0.0228). No acute effects were observed following treatment with CP101,606 or memantine.

In the protocol testing modulation of a negative affective bias 24hrs after treatment, the highest dose of HNK led to a positive affective bias in this test indicating re-learning with a more positive affective valence (one sample t-test: t11=2.916, p=0.0140, n=12) (Fig. 2A). HNK treatment at the lower dose of 1mg/kg (n=12, with Dunnett’s test, p<0.05, Fig. 2A) resulted in a sustained attenuation of the negative bias. The highest dose of lanicemine (repeated measures ANOVA, F3,33=6.099, p=0.0020, n=12, with Dunnett’s test, p<0.001, Fig. 2D) and both CP101,606 doses (1mg/kg: two-tailed paired t-test: t10=2.753, p=0.0204, n=11, Fig. 2E, and 3mg/kg: two-tailed paired t-test: t10=3.816, p=0.0034, n=11, Fig. 2E) as well as ephenidine (two-tailed paired t-test: t11=5.196, p=0.0003, n=12, with Dunnett’s test, p<0.001, Fig. 2F) also ameliorated the negative affective bias at the 24 hour timepoint, but did not induce a significant positive affective bias, although the p values for CP101,606 and ephenidine were less than p=0.1. PCP and memantine had no effect at 24hrs post treatment with all groups having similar negative affective biases.

**Figure 2:**
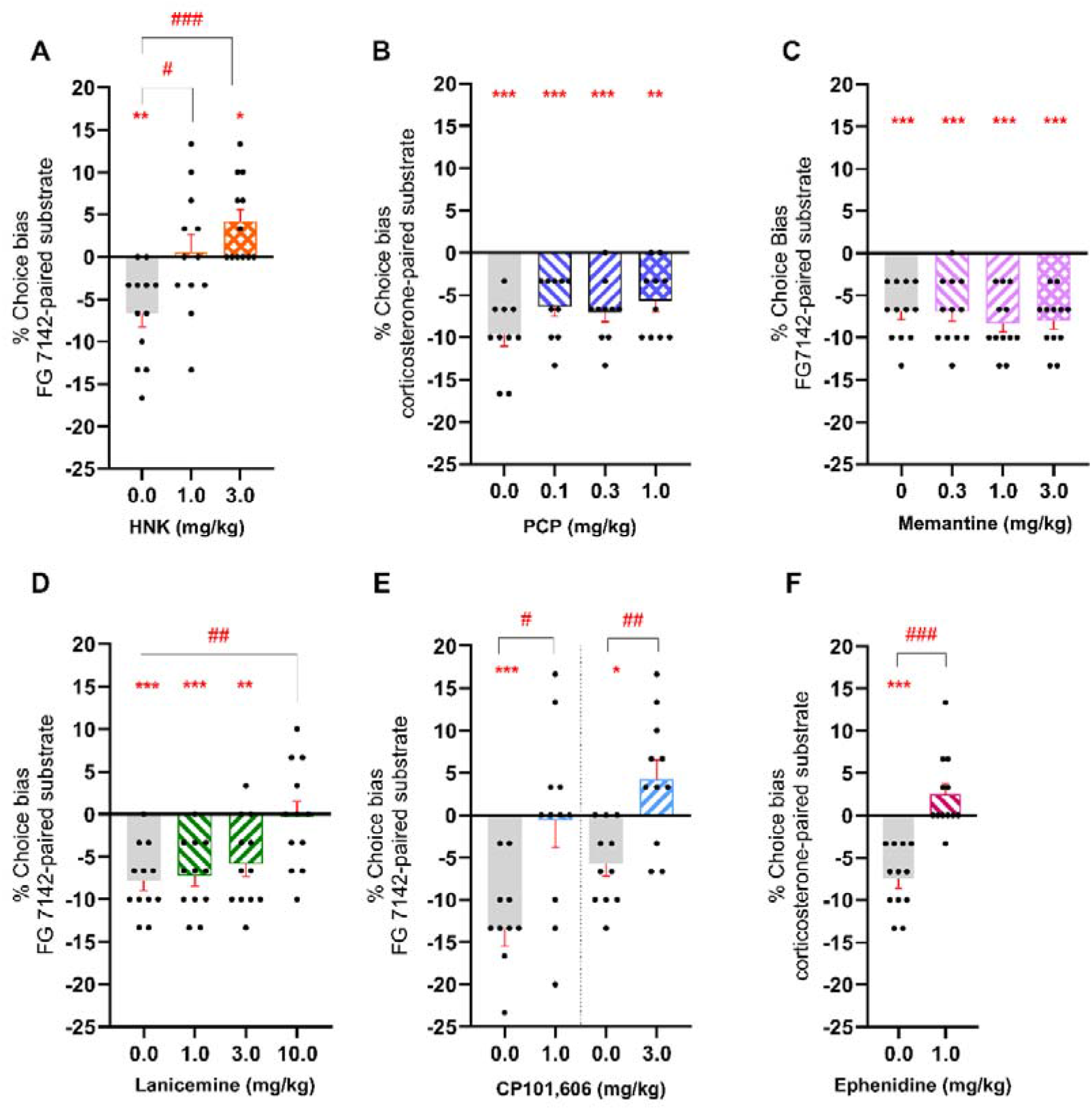
Treatment with HNK, lanicemine, CP101,606 and ephenidine at 24hrs prior to choice test resulted in sustained attenuation of negative affective bias in rats. Following FG7142- or corticosterone-induced negative bias rats were treated with HNK (0.3, 1.0, 3.0mg/kg; n=12; panel A), PCP (0.1, 0.3, 1.0mg/kg; n=10; panel B), memantine (0.3, 1.0, 3.0mg/kg; n=12; panel C), lanicemine (1.0, 3.0, 10.0mg/kg; n=12, panel D), CP101,606 (1.0, 3.0mg/kg; n=11, panel E) or ephenidine (1.0mg/kg; n=12, panel F) 24hrs before the choice test. Low dose of HNK 1.0mg/kg, the highest dose of lanicemine 10.0mg/kg, both doses of CP101,606 and ephenidine 1.0mg/kg significantly blocked the development of negative biases in rats, indication sustained attenuation similar to acute effects. Only the highest dose of HNK at 3.0mg/kg significantly inversed the bias into a positively valenced (panel A), similarly to the effects observed following ketamine treatment in Hinchcliffe et al. 2024. Data are shown as mean % choice bias ± SEM (bars) as well as individual data points (dots, n=10-12). Data were analyzed with one sample t-test against a null hypothesis mean of 0% choice bias (*p<0.05, **p<0.01, ***p<0.001) and pairwise comparisons were done using Dunnett’s test following main effect of treatment in RM ANOVA (#p<0.05, ##p<0.01, ###p<0.001).

Only the treatments where an acute modulation of affective bias had been observed were tested using the control, reward-learning assay. Animals treated with HNK, PCP, memantine, lanicemine and ephenidine developed similar reward-induced positive biases compared to the vehicle control, supporting specific affective bias modulation by the drug treatments (Figure 3A-E). No general memory impairments were observed.

**Figure 3:**
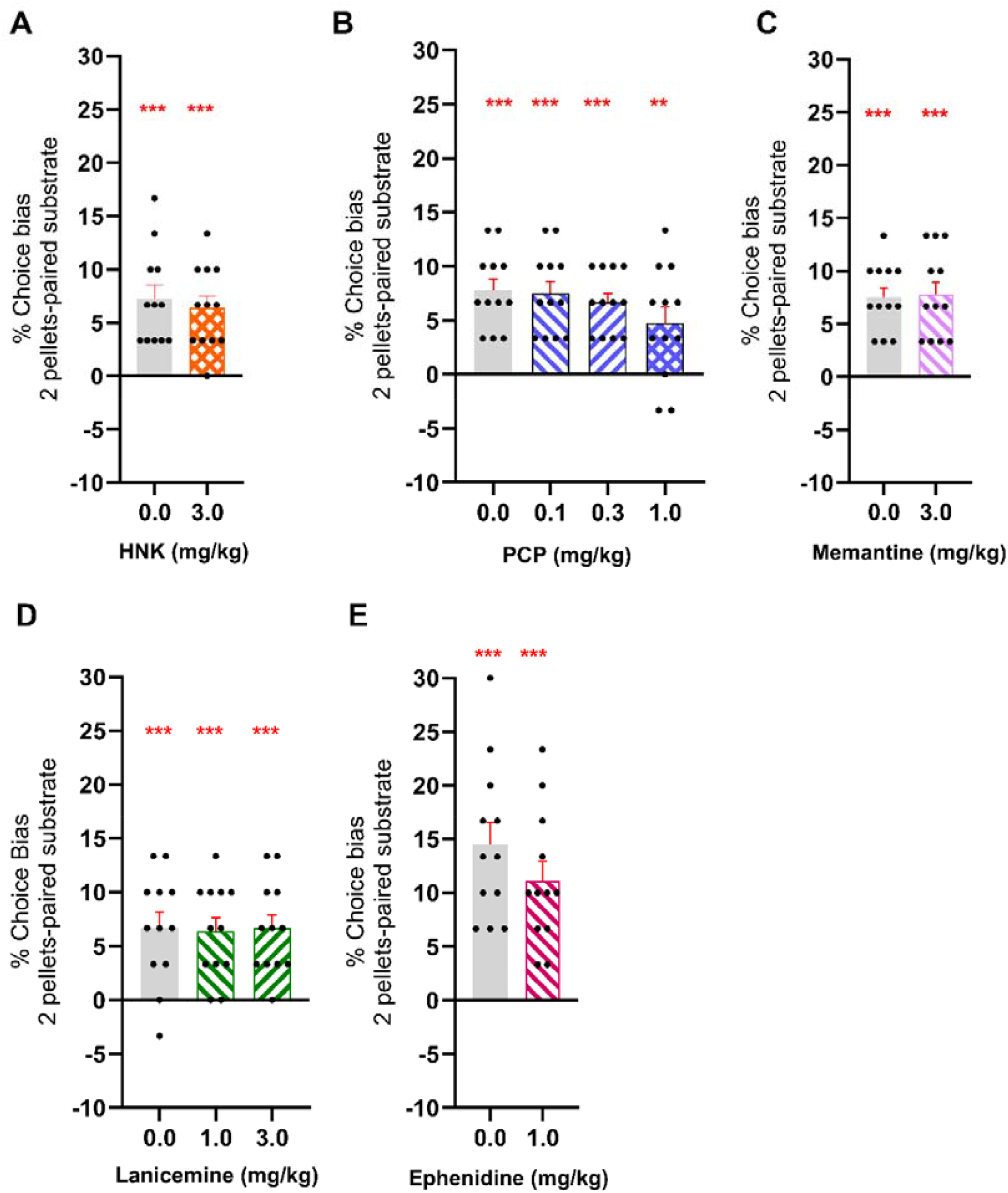
The reward learning assay has shown no general effects on memory retrieval. In the reward learning assay, a reward-induced positive bias was generated by using high (two pellets) versus low (one pellet) reward pairing sessions followed by administration of HNK (0.3, 1.0, 3.0mg/kg; n=12; panel A), PCP (0.1, 0.3, 1.0mg/kg; n=12; panel B), memantine (0.3, 1.0, 3.0mg/kg; n=12; panel C), lanicemine (1.0, 3.0, 10.0mg/kg; n=12, panel D) or ephenidine (1.0mg/kg; n=12, panel E) before the choice test. Data are shown as mean % choice bias ± SEM (bars) as well as individual data points (dots, n=12). Data were analyzed with one sample t-test against a null hypothesis mean of 0% choice bias (**p<0.01, ***p<0.001).

There was no evidence of non-specific impairments for the NMDAR antagonist treatments tested during choice tests (Tables S4, S6 and S8), apart from reward learning assay study with PCP treatment. Rats in the study with highest dose of PCP were significantly slower than vehicle group (Dunnett’s test, p<0.001). In a few studies changes in latency to dig and number of trials to criterion during learning pairing sessions were observed (for details, see Tables S3, S5 and S7). In the first week of acute effects of PCP, rats were slower to dig during pairing sessions with FG7142 comparing to the vehicle (paired t-test, t11=2.570, p=0.0261) and in the last week of sustained effects of lanicemine study, rats were faster to dig during pairing sessions with FG7142 (paired t-test, t11=5.124, p=0.0003). In the third week of acute effects of memantine study, rats did more trials to achieve criterion during pairing sessions with FG7142 (paired t-test, t14=2.987, p=0.0098).

## Discussion

Similar to the effects observed with ketamine, HNK, PCP, lanicemine and ephenidine acutely attenuated negative affective bias in rats with no effects observed with CP101,606 or memantine. When animals were tested 24hrs after treatment, of those having an acute effect, only HNK induced a positive affective bias similar to ketamine and suggesting modulation involving re-learning of the bias with a more positive affective valence. Lanicemine and ephenidine both demonstrated sustained modulated the retrieval of the negative bias but without clear evidence of a re-learning effect. Despite a lack of an acute effect, CP101,606 attenuated the negative bias at retrieval. Memantine had no effects in either assay. The lack of effects in the reward learning assay suggests any acute modulation observed was specific to affective bias modulation. The differences observed between the various NMDA antagonists tested suggest modulation of affective biases is sensitive to individual differences in their pharmacodynamic properties. Comparing the acute and sustained modulation of affective biases with these different compounds suggests the acute effects may be less predictive of antidepressant efficacy than the sustained effects. The lack of effects with CP101,606 at 1hr but sustained effects at 24hrs may relate to its lower acute dissociative effects and antidepressant effects respectively. Ephenidine is pharmacologically the most similar compound to ketamine and exhibits a similar profile of affective bias modification suggesting the NMDA receptor is the primary mediator of ketamine’s effects (19). Ephenidine lacks affinity for opioid receptor (19, 21) and does not produce the metabolite HNK. HNK’s effects were similar to ketamine but at doses which would be unlikely to be achieved from an effective dose of ketamine. HNK may have antidepressant effects but is unlikely to be the mediator of ketamine’s effects in terms of affective bias modification.

Our previous studies have shown that low dose of ketamine (1.0mg/kg) modulated biases in decision making and learning and memory (8, 14). In the judgement bias task (JBT), ketamine induced a positive change in cognitive bias index (CBI) following acute administration, indicating positive judgement bias (14). Ketamine has been extensively studied and at low doses its attenuation of negative affective biases, and the shift to more positive affective processing may involve synaptic plasticity in key brain regions (8). The re-learning effects of ketamine were dependent on protein synthesis, localised to the rat medial prefrontal cortex, and could be modulated by cue-reactivation, consistent with experience-dependent neural plasticity (8). Our current study expands upon these findings by demonstrating that other NMDA receptor antagonists, CP101,606 and ephenidine and HNK also produced sustained modulation of negative biases, albeit with varying efficacy and potency.

Consistent with findings in conventional models of antidepressant efficacy (11, 12), HNK attenuated negative biases at higher doses (1.0-3.0mg/kg) and the induced a positive affective bias 24hrs following treatment with the highest dose 3.0mg/kg. Since HNK requires a higher dose to achieve similar effects, it is unlikely that HNK is the sole reason why ketamine modulates affective biases at 1 mg/kg. Previous studies suggest HNK interacts directly with AMPA receptors to mediate its antidepressant effects (11, 23). These findings are important as HNK exhibits antidepressant-like properties similar to ketamine, however it lacks the psychotomimetic effects and abuse potential of its parent compound (11, 12).

Comparing different NMDAR antagonists, based on pharmacodynamic characteristics, our data suggests ketamine’s effects are not shared by other compounds. PCP is a potent very high ion trapping and non-selective NMDA receptor antagonist (24). In the affective bias test, PCP induced a robust attenuation of a negative biases acutely but failed to demonstrate any sustained changes on a negatively biased memory. Longer latencies following highest dose of PCP in the RLA indicate non-specific effects at higher receptor occupancy. These findings suggest the acute dissociative effects may have short term effects on the affective bias circuit but does not result in longer term modulation, or its more potent effects at the NMDA receptor limit specific modulation of the affective bias circuit.

The other NMDA antagonists tested, lanicemine and memantine, are low-trapping or partial trapping and moderate affinity respectively (23, 25, 26). Memantine had no effects while lanicemine had acute effects at all doses tested but only sustained effects at the highest dose of 10mg/kg. This dose likely generates much higher plasma levels and receptor occupancy than clinical doses and at more clinically relevant doses, no sustained effects were observed. The range of doses we tested covered the highest end of human dose range and higher. Memantine’s lower potency and partial trapping properties may explain its reduced efficacy compared to more potent antagonists such as ketamine. Both lanicemine and memantine failed to show antidepressant efficacy in clinical trials (23, 25, 26). This may be related to these differences in pharmacodynamic properties and suggests too high or too low potency reduces efficacy in terms of affective bias modification and may explain the reduced antidepressant efficacy observed.

To further explore this hypothesis, we tested ephenidine which is more similar to ketamine in terms of its pharmacodynamic properties. Ephenidine was more effective than the other NMDA antagonists tested with both acute and sustained modulation of affective biases observed at a similar dose to ketamine, 1mg/kg. Ephenidine blocks the NMDA receptor in a highly voltage-dependent manner (19, 27). Although these studies only looked at affective bias modification, the lack of affinity of ephenidine for opioid receptors suggests this is not the mediator of ketamine’s modulation of affective biases and suggests opioid receptors are not involved in ketamine’s antidepressant effects.

CP101,606 is a GluN2B subunit-selective NMDA receptor antagonist and has previously shown positive effects in a clinical trial in treatment resistant depression (17). The effects of CP101,606 were different from ketamine with no acute effects but a sustained modulation of affective biases. The sustained modulation of affective biases by CP101,606 suggests it is possible to dissociate between the acute and sustained modulation of affective biases and an acute effect is not necessary for the sustained effects to develop. The acute effects observed with the different NMDA antagonists tested suggest this may correspond with their acute dissociative effects, but these are not necessary for the antidepressant effects. CP101,606 has been reported to have less dissociative effects in patients and drugs specifically targeting NMDA receptors containing the NR2B subunit could provide better tolerated antidepressants.

## Conclusions

These findings provide important insights for understanding the role of NMDA antagonism in ketamine’s antidepressant effects and suggest pharmacodynamic properties may be important for efficacy. Findings in the affective bias test, and particularly the sustained modulation of negative affective biases, correspond well with clinical studies adding further to the hypothesis that modulation of affective biases is an important neuropsychological mechanism mediating antidepressant effects. Building from these studies, sustained affective bias modulation by novel NMDA antagonists in the affective bias test may provide a predictive model to identify more effective therapeutic interventions with reduced side effects and abuse liability.

## Author contributions

ESJR conceived and developed the methodology, and JKH contributed to the development and optimisation of methodology. JKH performed all statistical analysis, data visualisation and data curation. JKH conducted experiments investigating the acute effects of PCP, and sustained effects of PCP and ephenidine. JB conducted experiments evaluating the acute effects of HNK, CP101,606 and ephenidine, sustained effects of HNK and CP101,606 and the effects of HNK and ephenidine in the reward learning assay. KK conducted experiments investigating the acute and sustained effects of memantine and lanicemine and the effects of PCP, memantine and lanicemine in the reward learning assay. JB and KK contributed to the data analysis and data curation. JKH and ESJR wrote and edited the manuscript, and all authors reviewed and provided feedback on the manuscript.

## Data and materials availability

All data are available in the main text or the supplementary materials. Individual-level data for all studies are available on the Open Science Framework at DOI 10.17605/OSF.IO/MY8BN.

## References

1. Godlewska BR, Harmer CJ. Cognitive neuropsychological theory of antidepressant action: a modern-day approach to depression and its treatment. Psychopharmacology (Berl). 2021;238(5):1265–78.

2. Disner SG, Beevers CG, Haigh EA, Beck AT. Neural mechanisms of the cognitive model of depression. Nature reviews Neuroscience. 2011;12(8):467–77.

3. Roiser JP, Elliott R, Sahakian BJ. Cognitive mechanisms of treatment in depression. Neuropsychopharmacology : official publication of the American College of Neuropsychopharmacology. 2012;37(1):117–36.

4. Harmer CJ, Duman RS, Cowen PJ. How do antidepressants work? New perspectives for refining future treatment approaches. The lancet Psychiatry. 2017;4(5):409–18.

5. Hinchcliffe JK, Stuart SA, Mendl M, Robinson ESJ. Further validation of the affective bias test for predicting antidepressant and pro-depressant risk: effects of pharmacological and social manipulations in male and female rats. Psychopharmacology (Berl). 2017;234(20):3105–16.

6. Stuart SA, Butler P, Munafo MR, Nutt DJ, Robinson ES. A translational rodent assay of affective biases in depression and antidepressant therapy. Neuropsychopharmacology : official publication of the American College of Neuropsychopharmacology. 2013;38(9):1625–35.

7. Stuart SA, Butler P, Munafo MR, Nutt DJ, Robinson ESJ. Distinct Neuropsychological Mechanisms May Explain Delayed-Versus Rapid-Onset Antidepressant Efficacy. Neuropsychopharmacology : official publication of the American College of Neuropsychopharmacology. 2015;40(9):2165–74.

8. Hinchcliffe JK, Stuart SA, Wood CM, Bartlett JM, Kamenish K, Arban R, et al. Rapid-acting antidepressant drugs modulate affective bias in rats. Science Translational Medicine. 2024;16(729):eadi2403.

9. Berman RM, Cappiello A, Anand A, Oren DA, Heninger GR, Charney D, et al. Antidepressant effects of ketamine in depressed patients. Biological psychiatry. 2000;47(4):351–4.

10. Nikolin S, Rodgers A, Schwaab A, Bahji A, Zarate C, Jr., Vazquez G, et al. Ketamine for the treatment of major depression: a systematic review and meta-analysis. eClinicalMedicine. 2023;62.

11. Zanos P, Moaddel R, Morris PJ, Georgiou P, Fischell J, Elmer GI, et al. NMDAR inhibition-independent antidepressant actions of ketamine metabolites. Nature. 2016;533(7604):481–6.

12. Zanos P, Moaddel R, Morris PJ, Riggs LM, Highland JN, Georgiou P, et al. Ketamine and Ketamine Metabolite Pharmacology: Insights into Therapeutic Mechanisms. Pharmacol Rev. 2018;70(3):621–60.

13. Paoletti P, Bellone C, Zhou Q. NMDA receptor subunit diversity: impact on receptor properties, synaptic plasticity and disease. Nature Reviews Neuroscience. 2013;14(6):383–400.

14. Hales CA, Bartlett JM, Arban R, Hengerer B, Robinson ESJ. Role of the medial prefrontal cortex in the effects of rapid acting antidepressants on decision-making biases in rodents. Neuropsychopharmacology : official publication of the American College of Neuropsychopharmacology. 2020;45(13):2278–88.

15. Hinchcliffe JK, Robinson ESJ. The Affective Bias Test and Reward Learning Assay: Neuropsychological Models for Depression Research and Investigating Antidepressant Treatments in Rodents. Current Protocols. 2024;4(6):e1057.

16. Aguilar-Valles A, De Gregorio D, Matta-Camacho E, Eslamizade MJ, Khlaifia A, Skaleka A, et al. Antidepressant actions of ketamine engage cell-specific translation via eIF4E. Nature. 2021;590(7845):315–9.

17. Preskorn SH, Baker B, Kolluri S, Menniti FS, Krams M, Landen JW. An innovative design to establish proof of concept of the antidepressant effects of the NR2B subunit selective N-methyl-D-aspartate antagonist, CP-101,606, in patients with treatment-refractory major depressive disorder. J Clin Psychopharmacol. 2008;28(6):631–7.

18. Stuart SA, Wood CM, Robinson ESJ. Using the affective bias test to predict drug-induced negative affect: implications for drug safety. British journal of pharmacology. 2017;174(19):3200–10.

19. Kang H, Park P, Bortolotto ZA, Brandt SD, Colestock T, Wallach J, et al. Ephenidine: A new psychoactive agent with ketamine-like NMDA receptor antagonist properties. Neuropharmacology. 2017;112:144–9.

20. Stuart SA, Robinson ES. Reducing the stress of drug administration: implications for the 3Rs. Sci Rep. 2015;5:14288.

21. Morris H, Wallach J. From PCP to MXE: a comprehensive review of the non-medical use of dissociative drugs. Drug Testing and Analysis. 2014;6(7-8):614–32. https://www.3hs-initiative.co.uk

22. Zarate CA, Jr., Singh JB, Carlson PJ, Brutsche NE, Ameli R, Luckenbaugh DA, et al. A randomized trial of an N-methyl-D-aspartate antagonist in treatment-resistant major depression. Arch Gen Psychiatry. 2006;63(8):856–64.

23. Jentsch JD, Roth RH. The Neuropsychopharmacology of Phencyclidine: From NMDA Receptor Hypofunction to the Dopamine Hypothesis of Schizophrenia. Neuropsychopharmacology : official publication of the American College of Neuropsychopharmacology. 1999;20(3):201–25.

24. Sanacora G, Johnson MR, Khan A, Atkinson SD, Riesenberg RR, Schronen JP, et al. Adjunctive Lanicemine (AZD6765) in Patients with Major Depressive Disorder and History of Inadequate Response to Antidepressants: A Randomized, Placebo-Controlled Study. Neuropsychopharmacology : official publication of the American College of Neuropsychopharmacology. 2017;42(4):844–53.

25. Smith EG, Deligiannidis KM, Ulbricht CM, Landolin CS, Patel JK, Rothschild AJ. Antidepressant augmentation using the N-methyl-D-aspartate antagonist memantine: a randomized, double-blind, placebo-controlled trial. The Journal of clinical psychiatry. 2013;74(10):966–73.

26. Davies S, Alford S, Coan E, Lester R, Collingridge G. Ketamine blocks an NMDA receptor-mediated component of synaptic transmission in rat hippocampus in a voltage-dependent manner. Neuroscience letters. 1988;92(2):213–7.

