## Supplementary materials for "Modulation of negative affective biases in male rats may predict the rapid and sustained antidepressant effects of different NMDA antagonists"

| Cohort | Treatment | Dose (mg/kg) | Route of administration | Pre-treatment times |
| --- | --- | --- | --- | --- |
| 1, 2 | HNK | 0.0, 0.3, 1.0, 3.0 | IP (systemic) | 20min./24hrs |
| 3, 4 | PCP | 0.0, 0.1, 0.3, 1.0 | IP (systemic) | 40min./24hrs |
| 4, 5 | Memantine | 0.0, 0.3, 1.0, 3.0 | IP (systemic) | 60min./24hrs |
| 6 | Lanicemine | 0.0, 1.0, 3.0, 10.0 | IP (systemic) | 60min./24hrs |
| 2 | CP101,606 | 0.0, 1.0, 3.0 | IP (systemic) | 60min./24hrs |
| 1, 3, 7 | Ephenidine | 0.0, 1.0 | IP (systemic) | 60min./24hrs |
| 3, 7 | Corticosterone | 0.0, 10.0 | SC (systemic) | 30min. |
| 1, 2, 4, 5, 6 | FG7142 | 0.0, 3.0 | SC (systemic) | 30min. |

**Table S1: Summary of drug treatments in all animal cohorts.**

|  | Day 1 | Day 2 | Day 3 | Day 4 | Day 5 | Day 5/6 |
| --- | --- | --- | --- | --- | --- | --- |
|  | **Pairing 1** | **Pairing 2** | **Pairing 3** | **Pairing 4** | **Treatment with NMDAR antagonist** | **Choice Test** |
| Group 1 | CS+A vs. CS-  **Drug** | CS+B vs. CS-  **Vehicle** | CS+A vs. CS-  **Drug** | CS+B vs. CS-  **Vehicle** | **Drug A**  acute or 24hrs prior | CS+A vs. CS+B,  30 trials |
| Group 2 | CS+B vs. CS-  **Drug** | CS+A vs. CS-  **Vehicle** | CS+B vs. CS-  **Drug** | CS+A vs. CS-  **Vehicle** | **Drug B**  acute or 24hrs prior | CS+A vs. CS+B,  30 trials |
| Group 3 | CS+A vs. CS-  **Vehicle** | CS+B vs. CS-  **Drug** | CS+A vs. CS-  **Vehicle** | CS+B vs. CS-  **Drug** | **Drug C**  acute or 24hrs prior | CS+A vs. CS+B,  30 trials |
| Group 4 | CS+B vs. CS-  **Vehicle** | CS+A vs. CS-  **Drug** | CS+B vs. CS-  **Vehicle** | CS+A vs. CS-  **Drug** | **Drug D**  acute or 24hrs prior | CS+A vs. CS+B,  30 trials |

**Table S2A: Standard procedure for testing drug-induced affective bias versus vehicle.** Each animal receives drug treatment (Drug, i.e. corticosterone or FG7142) or vehicle counterbalanced over the four substrate-reward pairing sessions to induce a negative affective bias. On day 5 or 6 they undergo treatment with NMDAR antagonist (e.g. Drug A, B, C, D) either acutely or 24hrs prior to choice test. Substrate and day are also counter-balanced resulting in four different groups.

|  | Day 1 | Day 2 | Day 3 | Day 4 | Day 5 |
| --- | --- | --- | --- | --- | --- |
|  | **Pairing 1** | **Pairing 2** | **Pairing 3** | **Pairing 4** | **Choice Test with prior treatment with NMDAR antagonist** |
| Group 1 | CS+A vs. CS-  **2 pellets** | CS+B vs. CS-  **1 pellet** | CS+A vs. CS-  **2 pellets** | CS+B vs. CS-  **1 pellet** | **Drug A**  Choice test  CS+A vs. CS+B |
| Group 2 | CS+B vs. CS-  **2 pellets** | CS+A vs. CS-  **1 pellet** | CS+B vs. CS-  **2 pellets** | CS+A vs. CS-  **1 pellet** | **Drug B**  Choice test  CS+A vs. CS+B |
| Group 3 | CS+A vs. CS-  **1 pellet** | CS+B vs. CS-  **2 pellets** | CS+A vs. CS-  **1 pellet** | CS+B vs. CS-  **2 pellets** | **Drug C**  Choice test  CS+A vs. CS+B |
| Group 4 | CS+B vs. CS-  **1 pellet** | CS+A vs. CS-  **2 pellets** | CS+B vs. CS-  **1 pellet** | CS+A vs. CS-  **2 pellets** | **Drug D**  Choice test  CS+A vs. CS+B |

**Table S2B: Standard procedure for testing in the reward learning assay***.* Each animal receives 2 pellets or 1 pellet counterbalanced over the four substrate-reward pairing sessions, following the NMDAR antagonist treatment (e.g. Drug A, B, C, D) prior to choice test on day 5. Substrate and day are also counter-balanced resulting in four different groups.

| **Treatment** |  | **Response latency (s)** | | **Trials to criterion** | |
| --- | --- | --- | --- | --- | --- |
|  |  | **Vehicle** | **Drug** | **Vehicle** | **Drug** |
| Hydroxynorketamine | Week 1 | 3.3±0.4 | 3.2±0.3 | 6.5±0.2 | 6.7±0.3 |
|  | Week 2 | 3.1±0.4 | 3.3±0.5 | 6.3±0.2 | 6.1±0.1 |
|  | Week 3 | 3.4±0.5 | 3.1±0.3 | 6.5±0.3 | 6.7±0.3 |
|  | Week 4 | 2.8±0.4 | 2.7±0.3 | 6.4±0.3 | 6.0±0.0 |
| Ephenidine | Week 1 | 3.7±0.5 | 3.8±0.5 | 6.4±0.2 | 6.2±0.1 |
|  | Week 2 | 3.1±0.2 | 3.8±0.5 | 6.2±0.1 | 6.3±0.2 |
| PCP | Week 1 | **3.7±0.3** | **4.7±0.6***** | 6.8±0.1 | 6.7±0.1 |
|  | Week 2 | 2.7±0.1 | 2.8±0.2 | 6.3±0.1 | 6.3±0.1 |
|  | Week 3 | 2.4±0.1 | 2.4±0.1 | 6.3±0.1 | 6.2±0.1 |
|  | Week 4 | 2.3±0.1 | 2.3±0.2 | 6.3±0.1 | 6.3±0.1 |
| CP101,606 | Week 1 | 4.1±0.4 | 4.7±0.7 | 7.3±0.5 | 7.9±0.5 |
|  | Week 2 | 3.9±0.4 | 4.2±0.5 | 7.0±0.3 | 7.6±0.4 |
|  | Week 3 | 4.2±0.6 | 3.6±0.4 | 6.4±0.2 | 6.1±0.1 |
| Memantine | Week 1 | 1.6±0.1 | 2.0±0.0 | 6.3±0.2 | 6.3±0.2 |
|  | Week 2 | 1.8±0.1 | 1.9±0.1 | 6.3±0.2 | 6.8±0.2 |
|  | Week 3 | 1.8±0.1 | 2.1±0.1 | **6.8±0.3** | **8.0±0.3***** |
|  | Week 4 | 1.8±0.1 | 2.0±0.1 | 6.7±0.3 | 7.0±0.2 |
| Lanicemine | Week 1 | 1.5±0.0 | 1.6±0.2 | 8.5±0.7 | 9.3±0.6 |
|  | Week 2 | 1.6±0.1 | 1.6±0.0 | 7.5±0.5 | 7.1±0.4 |
|  | Week 3 | 1.6±0.1 | 1.6±0.1 | 7.8±0.5 | 7.0±0.3 |
|  | Week 4 | 1.4±0.1 | 1.4±0.1 | 7.1±0.4 | 7.2±0.3 |

**Table S3:** **Pairing sessions data: number of trials to criterion and latency to dig in the acute modulation studies**. Data shown as mean (n=11-12 animals/group) ± SEM averaged from the two pairing sessions for each substrate-reward association (vehicle/1 pellet or drug/2 pellets). There were no significant effects during pairing sessions, either on response latency to dig or number of trials to criterion following treatment with vehicle or any of the drugs. Only during first week of PCP study we observed difference in response latency (paired t-test, t11=2.570, p=0.0261) resulted in slower latency to dig during pairing sessions with FG7142 comparing to the vehicle and during third week of memantine we observed difference in trials to criterion (paired t-test, t14=2.987, p=0.0098) resulted in rats doing more trials to achieve criterion during pairing sessions with FG7142.

| **Treatment** | **Dose (mg/kg)** | **Response latency (s)** |
| --- | --- | --- |
| Hydroxynorketamine | 0.0 | 2.5±0.2 |
|  | 0.3 | 2.8±0.4 |
|  | 1.0 | 2.4±0.2 |
|  | 3.0 | 2.7±0.3 |
| Ephenidine | 0 | 3.1±0.3 |
|  | 1 | 3.0±0.4 |
| PCP | 0 | 2.4±0.2 |
|  | 0.1 | 1.9±0.1 |
|  | 0.3 | 2.2±0.2 |
|  | 1 | 1.8±0.1 |
| CP101,606 | 0 | 3.5 ± 0.3 |
|  | 1 | 2.9 ± 0.2 |
|  | 3 | 3.2 ± 0.4 |
| Memantine | 0 | 1.8±0.1 |
|  | 0.3 | 1.8±0.1 |
|  | 1 | 1.7±0.1 |
|  | 3 | 2.4±0.5 |
| Lanicemine | 0 | 1.4±0.1 |
|  | 1 | 1.4±0.1 |
|  | 3 | 1.5±0.1 |
|  | 10 | 1.5±0.1 |

**Table S4: Choice bias data: response latency to dig in the acute modulation studies.** Data shown as mean (n=11-12 animals/group) ± SEM averaged from the two pairing sessions for each substrate-reward association (vehicle or drug). No significant difference in latency to make choice was observed in studies following treatment with vehicle or any of the drugs: HNK, ephenidine, PCP, memantine and lanicemine.

| **Treatment** |  | **Response latency (s)** | | **Trials to criterion** | |
| --- | --- | --- | --- | --- | --- |
|  |  | **Vehicle** | **Drug** | **Vehicle** | **Drug** |
| Hydroxynorketamine | Week 1 | 3.4±0.4 | 3.7±0.4 | 6.6±0.2 | 7.2±0.4 |
|  | Week 2 | 3.7±0.3 | 3.6±0.3 | 6.6±0.2 | 7.1±0.4 |
|  | Week 3 | 3.0±0.2 | 2.9±0.2 | 6.8±0.2 | 6.5±0.3 |
| Ephenidine | Week 1 | 2.4±0.1 | 2.2±0.1 | 6.4±0.1 | 6.4±0.1 |
|  | Week 2 | 2.3±0.2 | 2.2±0.1 | 6.3±0.1 | 6.4±0.1 |
| PCP | Week 1 | 2.4±0.1 | 2.6±0.2 | 6.5±0.1 | 6.6±0.1 |
|  | Week 2 | 2.7±0.1 | 2.7±0.1 | 6.3±0.1 | 6.3±0.1 |
|  | Week 3 | 2.6±0.1 | 2.5±0.2 | 7.0±0.3 | 7.0±0.2 |
|  | Week 4 | 2.4±0.2 | 2.2±0.1 | 6.5±0.1 | 6.3±0.1 |
| CP101,606 1.0mg/kg | Week 1 | 3.7±0.5 | 3.9±0.6 | 6.0±0.0 | 6.0±0.0 |
|  | Week 2 | 4.6±0.6 | 4.9±0.7 | 6.7±0.3 | 7.0±0.3 |
| CP101,606 3.0mg/kg | Week 1 | 3.1±0.3 | 3.0±0.3 | 6.5±0.3 | 6.0±0.0 |
|  | Week 2 | 4.1±0.4 | 4.2±0.5 | 6.6±0.4 | 6.0±0.0 |
| Memantine | Week 1 | 1.5±0.0 | 1.5±0.0 | 8.6±0.5 | 7.5±0.2 |
|  | Week 2 | 1.6±0.0 | 1.5±0.1 | 8.4±0.5 | 7.2±0.3 |
|  | Week 3 | 1.5±0.0 | 1.5±0.1 | 8.8±0.4 | 7.6±0.2 |
|  | Week 4 | 1.5±0.0 | 1.5±0.1 | 8.6±0.6 | 7.4±0.3 |
| Lanicemine | Week 1 | 1.6±0.0 | 1.7±0.0 | 9.5±0.3 | 9.4±0.3 |
|  | Week 2 | 1.5±0.0 | 1.5±0.1 | 9.2±0.5 | 9.2±0.3 |
|  | Week 3 | 1.7±0.1 | 1.6±0.1 | 8.7±0.4 | 9.2±0.3 |
|  | Week 4 | **1.6±0.0** | **1.4±0.0***** | 8.1±0.2 | 8.9±0.3 |

**Table S5: Pairing sessions data: number of trials to criterion and latency to dig in the sustained modulation studies.** Data shown as mean (n=10-12 animals/group) ± SEM averaged from the two pairing sessions for each substrate-reward association (vehicle or drug). There were no significant effects during pairing sessions, either on response latency to dig or number of trials to criterion following treatment with vehicle or any of the drugs. Only during last week of lanicemine study we observed difference in response latency (paired t-test, t11=5.124, p=0.0003) resulted in faster latency to dig during pairing sessions with FG7142 comparing to the vehicle.

| **Treatment** | **Dose (mg/kg)** | **Response latency (s)** |
| --- | --- | --- |
| Hydroxynorketamine | 0.0 | 4.1 ± 0.5 |
|  | 1.0 | 3.9 ± 0.7 |
|  | 3.0 | 4.3 ± 0.7 |
| Ephenidine | 0 | 2.0±0.1 |
|  | 0.1 | 1.8±0.0 |
| PCP | 0 | 2.3±0.1 |
|  | 0.1 | 2.3±0.1 |
|  | 0.3 | 2.3±0.1 |
|  | 1 | 2.2±0.1 |
| CP101,606 | 0 | 3.8 ± 0.4 |
|  | 1 | 5.5 ± 1.1 |
|  | 0 | 3.9 ± 0.5 |
|  | 3 | 4.7 ± 1.1 |
| Memantine | 0 | 1.5±0.1 |
|  | 0.3 | 1.4±0.0 |
|  | 1 | 1.5±0.0 |
|  | 3 | 1.6±0.1 |
| Lanicemine | 0 | 1.6±0.1 |
|  | 1 | 1.6±0.1 |
|  | 3 | 1.6±0.1 |
|  | 10 | 1.5±0.1 |

**Table S6: Choice bias data: response latency to dig latency to dig in the sustained modulation studies.** Data shown as mean (n=10-12 animals/group) ± SEM averaged from the two pairing sessions for each substrate-reward association (vehicle or drug). No significant difference in latency to make choice was observed in studies following treatment with vehicle or any of the drugs: HNK, ephenidine, PCP, memantine and lanicemine.

| **Treatment** |  | **Response latency (s)** | | **Trials to criterion** | |
| --- | --- | --- | --- | --- | --- |
|  |  | **Vehicle** | **Drug** | **Vehicle** | **Drug** |
| Hydroxynorketamine | Week 1 | 4.0±0.5 | 3.3±0.3 | 6.5±0.2 | 6.7±0.2 |
|  | Week 2 | 3.5±0.3 | 3.1±0.2 | 6.5±0.2 | 6.4±0.2 |
| Ephenidine | Week 1 | 4.0±0.4 | 3.7±0.3 | 6.6±0.3 | 7.3±0.4 |
|  | Week 2 | 2.6±0.2 | 2.9±0.2 | 7.0±0.4 | 6.7±0.2 |
| PCP | Week 1 | 1.6±0.1 | 1.9±0.1 | 6.8±0.2 | 7.3±0.4 |
|  | Week 2 | 1.8±0.1 | 1.9±0.1 | 7.5±0.3 | 7.9±0.4 |
|  | Week 3 | 1.6±0.1 | 1.8±0.1 | 7.5±0.3 | 8.3±0.4 |
|  | Week 4 | 1.6±0.0 | 1.8±0.1 | 7.3±0.3 | 8.6±0.4 |
| Memantine | Week 1 | 1.5±0.0 | 1.6±0.1 | 7.8±0.2 | 7.6±0.3 |
|  | Week 2 | 1.5±0.0 | 1.5±0.1 | 6.9±0.2 | 7.8±0.5 |
| Lanicemine | Week 1 | **1.7±0.1** | **1.5±0.0***** | 9.3±0.4 | 8.3±0.3 |
|  | Week 2 | 1.6±0.0 | 1.5±0.0 | 9.3±0.4 | 8.4±0.3 |
|  | Week 3 | 1.7±0.1 | 1.7±0.0 | 8.3±0.3 | 7.9±0.2 |

**Table S7: Pairing sessions data: number of trials to criterion and latency to dig in the reward learning assay.** Data shown as mean (n=11-12 animals/group) ± SEM averaged from the two pairing sessions for each substrate-reward association (1 pellet or 2 pellets). There were no significant effects during pairing sessions, either on response latency to dig or number of trials to criterion following manipulation with 1 pellet versus 2 pellets. Only during first week of lanicemine study we observed difference in response latency (paired t-test, t11=2.499, p=0.0296) resulted in faster latency to dig during pairing sessions with FG7142 comparing to the vehicle.

.

| **Treatment** | **Dose (mg/kg)** | **Response latency (s)** |
| --- | --- | --- |
| Hydroxynorketamine | **0.0** | **3.5±0.3** |
|  | 3.0 | 3.2±0.2 |
| Ephenidine | 0 | 2.6±0.1 |
|  | 0.1 | 2.4±0.2 |
| PCP | 0 | **1.6±0.0** |
|  | 0.1 | **1.7±0.1** |
|  | 0.3 | **1.6±0.1** |
|  | 1 | **3.8±0.3***** |
| Memantine | 0 | 1.4±0.0 |
|  | 3 | 1.4±0.0 |
| Lanicemine | 0 | 1.5±0.1 |
|  | 1 | 1.5±0.0 |
|  | 3 | 1.5±0.0 |

**Table S8: Choice test data: response latency to dig in the reward learning assay.** Data shown as mean (n=11-12 animals/group) ± SEM averaged from the two pairing sessions for each substrate-reward association (1 pellet or 2 pellets). There were no significant effects during pairing sessions, either on response latency to dig or number of trials to criterion following manipulation with 1 pellet versus 2 pellets, in studies with HNK, ephenidine, memantine and lanicemine. The only significant difference was observed in the PCP (0.1-1.0mg/kg) study (RM ANOVA, F(3, 33) = 50.04, p<0.0001), rats were significantly slower to make a choice following highest dose (1mg/kg, p<0.0001) of PCP comparing to vehicle treatment.
